# Updated Transposable Element Libraries for *Drosophila melanogaster* in Dfam 4.0

**DOI:** 10.64898/2026.09.13.751287

**Authors:** Clément Goubert, Anthony Gray, Robert Hubley, Travis J. Wheeler, Arian A. F. Smit

## Abstract

*Drosophila melanogaster* repeatome, comprising roughly 20% of the genome, is characterized by a large fraction of active TE families counterbalanced by efficient purifying selection. Consequently, many TE families persist at low copy numbers and are frequently population specific. Hybridization and horizontal transfer provide new families, which can spread through natural populations within decades. These dynamics, together with years of independent curation efforts, left the *D. melanogaster* mobilome distributed across several, partly redundant, libraries. Prompted by submissions of population-specific data, we undertook a complete overhaul of the *D. melanogaster* TE libraries, begun in Dfam 3.9 and finalized in Dfam 4.0. We cross-referenced the new submissions against Repbase, FlyBase, the Berkeley Drosophila Genome Project, and our own Dfam 3.8 to resolve redundancy and reconcile names, then rebuilt or newly constructed the seed alignment for most families. Seeds came from four sources: the dm6 reference itself, which supported the majority of models; insertions >100 bp from 13 samples of a recently published *D. melanogaster* pangenome; the genomes of other members of the *D. melanogaster* subgroup, which supplied copies for older families too degraded in dm6 alone; and diverged matches recovered during iterative curation, which resolved into subfamilies and previously undescribed relatives. Rebuilding the seeds corrected consensus sequences that were truncated, chimeric, or skewed by co-duplicated fragments, and lowered the mean Kimura divergence of annotated copies from their consensus. The revision also added families with no prior Dfam representation, including the DNA P-element (absent from dm6) and a collection of novel families that have recently invaded natural populations. Following manual curation, the new library contains 398 models, up from 226 in Dfam 3.8, and annotates an additional 1.5% of the dm6 reference.

## Background

The fruit fly *Drosophila melanogaster* is a foundational model in genetics. Its short generation time, large brood sizes, easily trackable phenotypes, and giant polytene chromosomes, which enable the direct observation of gene activity, have made it a powerful experimental system^1^. Arguably, *D. melanogaster* is one of the best studied animal genomes to date, reflecting more than a century of scientific investigation^2^. Amid rapid improvement of molecular biology techniques and a renewed interest in Barbara McClintock’s work on controlling elements, the first mobile elements in the genome of *D. melanogaster* were identified in the late 1970s, laying the foundation for decades of research into their biology and evolution^3^.

Unlike most mammalian genomes, where TE copies typically account for >50% of the sequence and few active families persist, in fruit flies TE copies are short lived owing to high genomic deletion rates^4^ and efficient purifying selection^5,6^. TE copies in *D. melanogaster* stem from families whose activity vary widely between populations^7–10^. While the dm6 reference genome has an estimated interspersed repeat content of ∼15% in an assembly size of 137 Mb (ignoring ambiguous (N) nucleotides), the actual TE content is estimated to range between 16 and 21%^8^, with some TE families absent from some populations. For instance, the P-element, which has been abundantly studied for causing hybrid dysgenesis^11^, is absent from the reference genome dm6 which was derived from a lab strain isolated from wild populations. Notably, novel TE families have been shown to be able to spread worldwide within a few decades after a horizontal transfer event, a process aided by human activity^12,13^.

Widespread interest in the model, combined with scattered representation of the *D. melanogaster* mobilome across multiple populations, has led to a proliferation of TE libraries. The resulting collection of partially redundant and uncoordinated databases of TEs has been a source of confusion and inconsistency in the literature. Historically, several respected projects have curated and maintained databases of consensus sequences for *D. melanogaster*, including Repbase^14^, the Berkeley Drosophila Genome Project (BDGP)^15^ and the Flybase consortium^16^. In 2015, in collaboration with Repbase, Dfam developed a library of seed alignments and profile hidden Markov models (pHMMs) for *D. melanogaster* TE families^17^. Dfam’s models were created by building seed alignments from genomic instances identified by RepeatMasker from the dm6 reference genome, using the Repbase consensus sequences.

Through the years, new research has led to the identification of additional elements in *D. melanogaster* genomes^12,13,18^, the most comprehensive effort stemming from the *de novo* construction of a manually curated library (MCTE) based on 13 long read assemblies of diverse populations^10^. In 2023, the MCTE library, as well as a handful of population specific elements identified by Pianezza et al. (2025) were deposited to Dfam and prompted us to carefully reevaluate the standing of the TE library available for *D. melanogaster*.

### Desirable Traits for TE libraries

In developing this library and descriptive manuscript, we have found it useful to explicitly define the characteristics that we desire in a library of TE families. We describe a few core principles here, so that they are documented.

### What makes an ideal TE library?

From a biological perspective an ideal library of models for a given species would have high-quality models, including seed alignments and pHMMs, for all TE families that have left remnants in its genome.

In reality, low copy-number TEs may not be perfectly reconstructed due to regional coverage limitations. For ancient copies, the accumulated mutations may make a seed alignment inaccurate. Given enough time, TE copies will not even be recognized as interspersed repeats. Even with many relatively young copies, class I TEs rarely can be reconstructed precisely. Usually, the consensus represents an artificial average of a population of evolving class I TEs, given their often-rapid evolution in a genome and the parallel activity of multiple source genes. These evolutionary patterns can, to a limited extent, be expressed by the creation of multiple subfamily models^19,20^, which can be achieved today using our software coseg^21^ and other scripts discussed in Storer et al.^22^.

In practice, libraries for species are used for genome annotation with competitive search software such as RepeatMasker, Censor^23^, or TEannot^24^. A good library for this purpose would have the following characteristics:

- The library should be devoid of redundancy to avoid collision between annotations, maximizing reproducibility.
- It should not contain models merely representing low complexity DNA, which are commonly created by de novo library building software. These models are a key source of false annotations, because all sequences in the genome with similar composition will produce alignments to such models with inappropriately-high scores, simply due to the shared composition of the sequences.
- The library should not contain reconstructions of cellular genes, in order to avoid annotating those genes as TEs. In mammals and other species with LINE1 activity, a common cause of this problem are high copy number processed pseudogenes, which manifest as interspersed repeats.
- In our experience, homology searches benefit from representing the LTR regions and internal sequences of LTR retrotransposons with separate models. With the same classification and the suffixes _LTR and _I, RepeatMasker will indicate flanking matches to LTR and internal models as a single TE insertion by giving the matches the same ID.
- A library of models for a given species does not benefit from the unlimited expansion of subfamily models as more and more copies cannot be unambiguously mapped to one particular model^25^. A common practice is to exclude models that are >80% similar over >80% of their length to an existing model. In practice, these numbers are too severe and lead to diminished sensitivity for the detection of older elements and of unrelated or highly diverged regions of the subfamilies, as well as an overestimate of the age of the excluded TE copies based on the substitution level from the chosen consensus. For subfamilies that can be aligned end-to-end it is possible to make a combined seed alignment in which the subtler subfamily structure can still be acknowledged and displayed by grouping the seeds in the seed alignment and/or inclusion of a tree of the seeds’ relationship.
- The classification given to the model should follow a recognized classification system. Dfam has implemented a trackable system that combines concepts from established systems with phylogenies based on reverse transcriptase and transposases (see also: https://dfam.org/classification/tree)
- Each model has a taxonomic label which indicates the highest taxonomic group where all members share orthologous copies of the TE. In case of horizontal transfer there will be multiple clade names. Dfam will include the model for all species belonging to these clades
- Recognition should be given to the source of the model, i.e. the curator or consortium responsible for building the seed alignment for each family.
- A minimal description of the library building process should be available.
- A reference to the first publication describing the TE family should be associated as metadata
- If known, aliases to different names given for the same element in other databases should be provided

### Seed alignments as foundation of TE libraries

As introduced in the background section, Dfam’s TE family models are built on the crucial concept of seed alignment, where seeds are selected to represent the richness and diversity of sequences of its members. An ideal seed alignment has the following characteristics:

As most TE copies decay in a neutral fashion, the sequences included in the seed alignments should show a relatively narrow, random distribution of substitution levels as expected by chance. High divergence outliers generally indicate the presence of copies of a related TE and should be removed (and ideally be used to create their own model(s)). A subset of copies that are much more similar to each other than to the consensus often have been recently duplicated via other mechanisms, for example via segmental duplications or satellite expansion, and can distort the consensus and model. They are usually distinguished by shared flanking DNA and should be reduced to one member. Note that functionally adopted (exapted) TE copies may be conserved outliers as well, but do not share conservation outside the TE-derived sequence.
The per position coverage over the length of the model should be regular. Dependent on the biology of the element, this coverage need not necessarily be uniform; for example LINE models show a gradually increased coverage towards the 3’ end (due to the random 5’ truncation at insertion) and many class II transposons show blocks of higher coverage at the termini, due to the transposability and proliferation of internal deletion products. In contrast, many automatically created models show a very irregular coverage pattern. Short regions of relatively very high coverage in a seed alignment often indicate the (partial) alignment against copies of distinct more abundant TEs; this may occur when the other family has regional homology or when a copy of that family was naturally embedded in the modeled TE. Another common pattern is the presence of two or more regions of abruptly different coverage caused by the existence of TEs with sequence similarity through recombination events. In both cases, the fragmentarily aligned seeds should be removed as they distort the true nature of the modeled TE.
Unless extant copies are still near-identical to their source genes, coverage should be at least three-fold over the length of the model. Low coverage can often be overcome by collecting seeds from related species or from pangenomes.
Target sites duplicated during integration, e.g. TA for the Tc1/Mariner group of transposons, should not be part of the model. However, it is useful to include the type (and frequency) of observed target site duplications as metadata, as it supports classification and confirms that a model is complete and does not represent a chimaera or hybrid.
The consensus sequence derived from a seed alignment should represent the full-length, precise reconstruction of the active transposable element (source gene)

○ The orientation is that of the main (e.g. transposase) coding region. Non-autonomous elements without a coding region should be in their transcriptional orientation for class I (retrotransposing) elements and others in that of known autonomous elements sharing sequence similarity.
○ All coding regions of (functional) genes have full open reading frames and, where present, intact and in-frame splice sites. Note though that sometimes seemingly autonomous elements have defective coding regions and depend on other copies for transposition.

## Methods

Motivated by the submission of the MCTE library to Dfam, we started the revision process by comparing the MCTE library with Dfam 3.8 entries, removing identical or near-identical sequences (see *Elimination of near duplicates)*, and prioritizing full-length TE consensus entries. In addition, we considered a handful of elements that recently swept through novel populations^12^, also submitted to Dfam, and the consensus sequences discovered in *D. melanogaster* by Bargues and Lerat^18^. Each consensus was cross-referenced against Dfam and other existing libraries (see *Gathering of additional models from existing databases*) to identify missing entries and establish aliases and relationships. For families absent from Dfam, new seed alignments were created using *D. melanogaster* dm6 when sufficient copies existed to create a high-quality seed alignment (see *Model building*); otherwise, an augmented dm6 assembly incorporating insertions from 13 alternative genomes^10^ was used (see *Pangenome search of candidate seed alignments)*. All new and existing TE family models underwent manual review, with particular focus on seed alignment support, leading to many model refinements.

Each TE model was further annotated with ORFs, target site duplications (TSDs), and in the case of LTR elements, splitting into LTR and internal (INT) regions following a thorough manual curation process. Together, these steps improved annotation accuracy and completeness, resolved naming ambiguity across databases and yielded novel, well-supported entries.

### Strategies for model improvement and expansion

We employed a number of parallel strategies to improve and expand the existing Dfam library for Drosophila melanogaster.

Dfam 3.8 contained 238 models which in 2015 had been automatically derived from copies identified by a RepeatMasker run on dm6 with the *D. melanogaster* consensus sequences present in Repbase at that time (release 20.07); of these, 12 were later found not to be interspersed repeats, leaving the 226 models that we compare against below. This included 9 models for satellite repeats, 1 for 5S RNA, 12 mostly unclassified elements that we found not to be interspersed repetitive nor TE-derived, and 3 models for TEs not found in *D. melanogaster* (the LTR and internal sequence of TOM from *Drosophila ananassae* and the Tc1-like DNA transposon UHU from *Drosophila heteroneura*). Given the presence of 69 LTR and internal pairs, the remaining 213 models represented 144 different TEs.

Many of the seed alignments for these models were of such low quality that the consensus sequences derived from them were significantly diverged from the original Repbase input. In the worst cases, large regions were missing due to a dearth of copies in dm6. Many seed alignments were contaminated with cross matches to other TEs or warped by a large number of fragments that had co-duplicated inside a tandem repeat. Other TEs were incomplete because the original Repbase input was fragmentary. We rebuilt seed alignments for 176 models that either showed problematic seed coverage, produced a consensus differing by more than 5% from the original Repbase input or appeared to be incomplete.

### Gathering of additional models from existing databases

Besides exploring the MCTE library, we collected consensus sequences for TE families present in *D. melanogaster* from the following publicly accessible databases: Repbase 27.06 (last public release), Flybase (FB2023_05 September, Dmel Release 6.54) and BDGP^26^,*

We compared these libraries to Dfam and each other to find unrepresented elements. In order for an entry to be considered potentially new, it should match a number of interspersed copies in a competitive search with the combined databases against the fruit fly pangenome. For example, the MCTE library had 165 entries, 52 of which did not correspond to a Dfam entry. However, 18 of these did not represent *D. melanogaster* TEs: 16 were fragments of local near-tandem duplications, one was a fragment of the mitochondrial genome, and one was a small fragment of the *D. virilis* Penelope element^27^, which has not yet been reported in *D. melanogaster* populations though it has been artificially introduced in the lab in the *D. melanogaster* genome^28^. Besides the 34 new consensus sequences from MCTE, we found 13 in Flybase and 11 in BDGP. Also, the most recent open Repbase version available at that time (27.06) had 83 additional *D. melanogaster* models (for 43 TEs and 3 satellites). Some of the elements were shared between the database and many were fragmentary. At the end we could build 103 full-length new models for 59 different TEs (44 LTR elements were separated in 2 models) based on this input.

### Finding new elements similar to those in closely related species

Due to both high frequency of introgression^29^ and recent speciation times, many TEs have been active in the genomes of closely related Drosophila species or in that of a common ancestor. The latter may lead to the presence of distinct subfamilies in each lineage. We therefore also scanned the *D. melanogaster* genome with all Repbase 27.06 entries for TEs in the other members of the *D. melanogaster* species subgroup (*D. simulans*, *D. mauritania*, *D. sechellia*, *D. yakuba* & *D. erecta*). We could build another 24 models for 13 TEs using this approach. 17 of these models were different enough from the query to warrant a new name for the element found in *D. melanogaster*.

### Finding new elements distantly matching known D. melanogaster TEs

While retrieving copies for a TE from the (pan)genome, one often recovers significant matches that are much more diverged from the consensus than the copies of said TE. We collected such matches and, if not corresponding to another known TE, built seed alignments for them using iterative searches with improved consensus sequences to gather more and longer seeds. Some of the new models constituted subfamilies of the original query, but more often the similarity was distant, and new families could be described.

With the last two approaches we created 73 models for 2 satellites and 40 TEs (31 LTR elements) that were not yet in any of the input databases.

### Model building

Following comparison between novel entries and existing databases, many TE families were represented in more than one collection and some TEs had redundant entries in the same database. We clustered the elements and worked on each cluster individually. Usually, clustered elements represent the same TE, but they may also reflect true subfamilies. Using the prioritized Dfam entries, or the representative of a novel submission (non-redundant with Dfam), we first collected all copies from the reference dm6 genome with a cutoff score above 250 using RepeatMasker parameters, capturing flanking DNA to observe target site duplication or allow expansion of incomplete models or only partially detected subfamily copies. When insufficient seeds were retrieved for a seed alignment, additional seeds were gathered: for young elements, additional seeds were taken from a pangenome (see below), and for older elements, seeds were taken from the genomes of other members of the *D. melanogaster* species group. For example, we used *D. sechellia* to supplement the seed for Invader1_I, (DF000001649). In each case, the relevant taxonomic level was also automatically assigned (e.g., for Invader: the melanogaster subgroup).

Near-identical copies produced by post-transposition duplication were collapsed to a single seed instance (<u>DeleteAlmostDupfromSeqfile.pl</u>) to keep them from biasing the models. This is a critical step for a genome, like that of *Drosophila*, with many relatively low copy number repeats. This problem is aggravated by the seemingly high number of local duplications of TE fragments, perhaps correlated with the location of (old) piRNA clusters. We used the scripts alignAndCallConsensus.pl to build and, if necessary, extend models to their full length. During this process, we compare the seeds to the current model and also to all related or partially matching database elements, which function as “buffers” to avoid incorporating their copies in the seed alignment. We used the script AutoRunBlocker.pl to resolve gaps and poorly aligned subregions almost always to the point that the derived consensus sequence has full open reading frames and termini characteristic for their class. Usually, the full extent of the TE could be confirmed by observing type-specific target site duplications flanking one or more copies. Issues can arise when tandem repeats are found in TE family models. For example, the foldback DNA transposon NOF_FB (DF000001677) carries a satellite-like tandem repeat embedded within its long terminal inverted repeats (TIRs), which causes alignment tools to collapse the two near-identical TIRs onto a single copy in the consensus (the 5_′_ TIR absorbs the matches while the 3_′_ TIR shrinks away). This was resolved by aligning the copies twice against the consensus, once with the 5_′_ and once with the 3_′_ TIR masked, so that each run could only match one TIR at a time and the two ends were no longer confused; a low cross_match masklevel of 80%, together with a reduced bandwidth (-ba 5) and a high gap penalty (40) further kept the satellite from producing spurious, over-extended alignments.

### Break-down of LTR elements

The boundaries between the long terminal repeats (subsequently labelled with the suffix “LTR”) and the internal coding region (“INT” or “I’) of each curated LTR retrotransposon family were defined from a combination of structural features rather than any single signal. The breakpoints of the LTR regions are expected to match the two identical sub-hits, aligned to the 5’ and 3’ end of the query, produced by self-comparison of the family consensus; however, automatic reconstruction of these elements sometimes produces truncated LTRs flanking an (overextended) internal region. Complete terminal repeats are expected to fall within a length range of roughly 180-2000 bp and to carry the canonical 5_′_-TG…CA-3_′_ ends motifs, or, for 72 of the 133 *D. melanogaster* Gypsy elements, a non-canonical 5_′_-AGTNA…TAANT-3_′_ terminus. A polyadenylation signal toward the 3_′_ LTR served as an additional landmark. The internal edge of the 5_′_ LTR was further constrained by locating the primer-binding site (PBS), the short tract immediately 3_′_ of the 5_′_ LTR that is complementary to the 3_′_ end of a host tRNA. Mature tRNA sequences (truncated to their final ∼30 nt, as PBSs match the final 8-25 tRNA nucleotides) were obtained from GtRNAdb^30^ (https://gtrnadb.ucsc.edu/), with the 3_′_ CCA appended manually (GtRNAdb sequences omit it) and aligned against the consensus; the *D. melanogaster* tRNA set was supplemented with vertebrate tRNAs to cover tRNA types potentially missing from current annotations. PBS matches were typically found within ∼35 bp of the INT start and most often immediately adjacent to, or overlapping by one base, the 3_′_ end of the 5_′_ LTR. Alignments used cross_match-based comparison with the 14p49g scoring matrix distributed with RepeatMasker and a masklevel setting >= 98, the latter to prevent a genuine terminal tRNA match from being displaced by a spurious higher-scoring overlap. For families with detectable similarity to previously characterized elements, LTR and INT segments were additionally assigned by comparison to the Dfam database.

Repbase and Dfam have separate LTR and INT models because of the high frequency of solitary LTRs in some organisms; in mammals, for example, many LTR elements are only known by their LTR. Other reasons to maintain separate models are (i) the faster evolution of the LTRs and (ii) the propensity for LTR element transcripts to recombine during the reverse transcription process. When few copies are retained, as is often the case in Drosophila, establishing models for LTR elements becomes hazardous. For example, the original Stalker element has only 2 copies in the genome and its LTRs are usually found flanking Stalker4, while the Stalker3 internal sequence is sometimes found with Stalker2 LTRs (the recombination point in either case happened to be near the INT/LTR boundary).

### Pangenome search of candidate seed alignments

In order to build robust seed alignments for new models, including families with low or no support in the reference genome dm6, we built an augmented reference by adding structural variants absent from the dm6 reference but present in the 13 genome assemblies used by ^10^ to build the MCTE library. We used the first stage of GraffiTE^31^ which implements the minimap2/SVIM-asm^32,33^ pipeline to detect all insertions >100 bp. All alternative alleles (insertions relative to dm6) were extracted from the VCF output and concatenated to the dm6 assembly in FASTA format. Next, candidate consensus sequences were searched against the augmented dm6 assembly with Repeatmasker version 4.1.6, using the RMblastn search engine in slow search mode (-s). We finally generated new seed alignments from the Repeatmasker output using the script generateSeedAlignments.pl distributed with Repeatmodeler2^34^.

### Comparing Dfam 4.0 to all contributing libraries

To establish a measure of similarity between consensus sequences in the final Dfam 4.0 library and sequences in each contributing database (Dfam 3.8, Repbase, Flybase, BDGP and MCTE), we computed a score based on pairwise alignments between sequences. Alignments were performed with cross_match wrapped by our Xmatch.pl in -perbasescore mode (see Availability of Data and Materials). This mode produces a similarity measure that is more nuanced than simple percent identity, comparing the alignment score of the two sequences with the maximum possible score. Specifically: using a symmetric nucleotide scoring matrix, (i) the denominator is computed as the maximum possible score, which is the score of a self-alignment of the Dfam 4.0 consensus (the sum of per-position match scores across the length of the sequence, using lowest self-score for any ambiguous bases (N and IUPAC codes)), and (ii) the numerator is computed as the raw (non-complexity-adjusted) alignment score for the Dfam 4.0 consensus against the other sequence.

### Reference genome annotation

The reference genome dm6 was downloaded from the UCSC genome browser^35^*. Drosophila melanogaster* TE libraries were extracted in FASTA format from Dfam versions 3.8 and 4.0 using the python package FamDB (accessible online at https://github.com/Dfam-consortium/FamDB) with the following command:

`./famdb.py families -ad “Drosophila melanogaster” --curated -- format fasta_name --include-class-in-name > library.fa.`

Repeatmasker v.4.2.3 was then run on the reference genome using RMblastn as its search engine with the command: `RepeatMasker -a -lib library.fa dm6.fa`. Summary statistics from Repeatmasker runs, including binned Kimura substitution levels for the TE landscape analysis, were generated with the utility scripts buildSummary.pl and calcDivergenceFramAlign.pl distributed with Repeatmasker.

## Results and Discussion

### Families added and removed

For *Drosophila melanogaster*, the curated section of Dfam 4.0 now contains 398 models, including 371 TEs, 11 satellites, 7 rRNAs and 9 sequencing artefacts models, nearly doubling the total in Dfam 3.8 of 226 models. The current release also contains seven additional models that were flagged after the 4.0 release as artefacts, non-repetitive or erroneously assigned to Metazoa and will be removed from Dfam with the next release (their effect on annotation is negligible; Table S1). The final count of models does not directly reflect the number of distinct TE families, as the LTR (N = 152) and internal sequences (N = 130) of LTR retrotransposons are represented by different models. We created a total of 303 new or improved seed alignments, while 87 seed alignments in Dfam 3.8 were unchanged.

These improvements had several origins. (1) New TE families absent from Dfam were contributed by researchers mining for young TEs in population level data. For example, the MCTE library^10^, which was built from 13 non-reference genomes sourced from wild strains added novel population-specific families, while^12^ provided some families, such as Spoink, Souslik, Micropia2 and Transib1Y, that have recently invaded natural populations (Figure 1). (2) We surveyed legacy databases (Flybase, BDGP, the public Repbase) and also explored if TEs observed in other *Drosophila* species may have relatives that left copies in the *D. melanogaster* genome. (3) While improving seed alignments, more diverged matches occasionally could be clustered to create a model for a new subfamily or even a (distantly) related TE. (4) For older elements represented by few and fragmentary copies in the *D. melanogaster* genome we sometimes could complement the seed alignments with (preferentially non-orthologous) copies in the genomes of other members of the *D. melanogaster* species subgroup (*D. simulans, D. maurtiana, D. sechellia, D. yakuba & D. erecta*). Such a case is illustrated with the Gypsy element Invader5 (Figure 2). (5) For very young elements with poor representation in the reference genome dm6, we could improve or complete models by collecting copies from an augmented reference genome, containing all insertion sequences (>100bp) present in the 13 genomes of ^10^.

**Figure 1.**
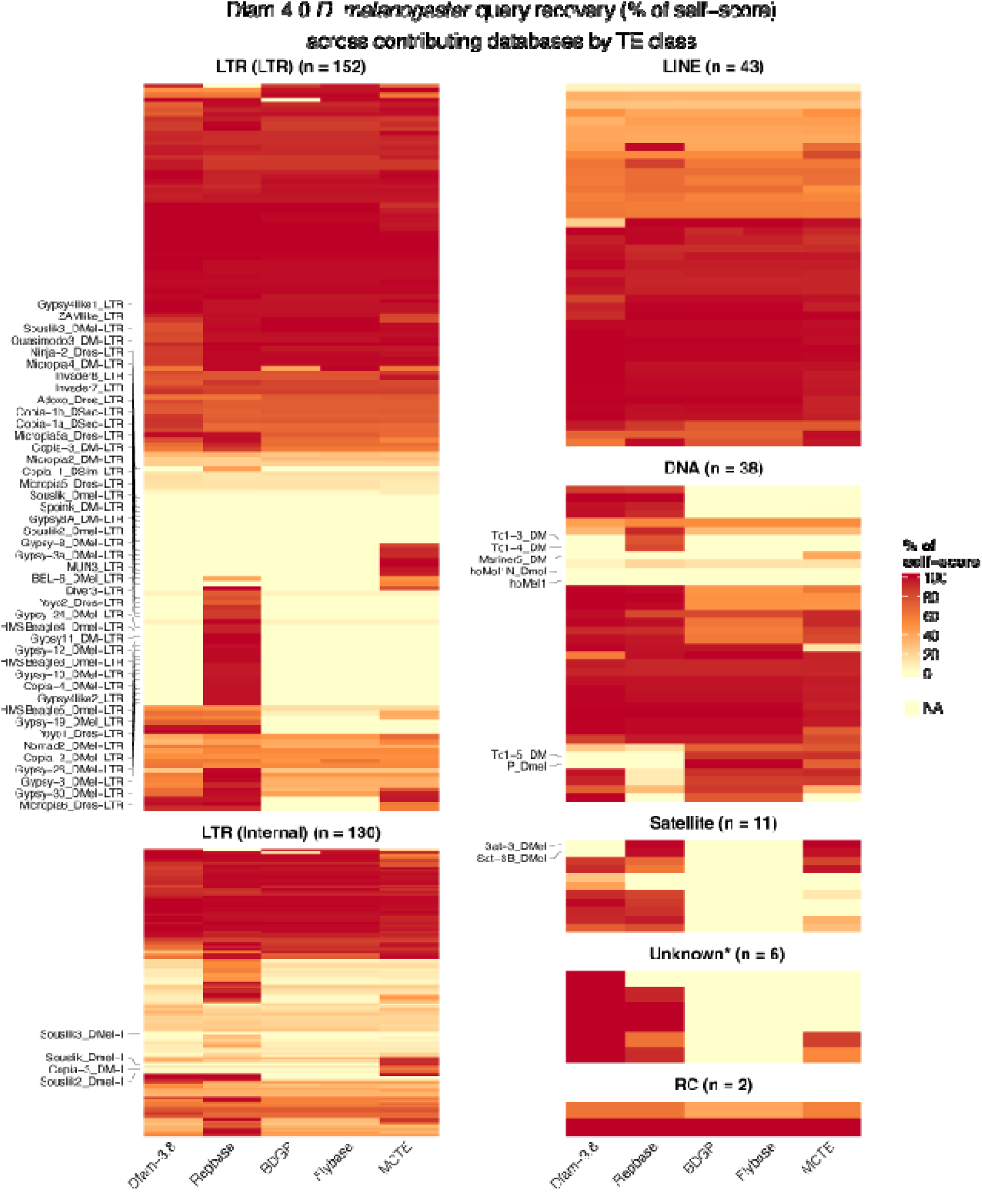
Contributing library relationships to Dfam 4.0. In these heatmaps, each row corresponds to a model in the Dfam 4.0 *D. melanogaster* library. Each column corresponds to one of the contributing libraries (Dfam 3.8, Repbase, etc). The color of a cell indicates the similarity (in percent of self-score, see Methods) between the 4.0 model and the most similar model found in the column’s library, from 100 (dark red, sequences effectively identical) down to the lowest-scoring detectable match (yellow). Cells are left blank where the Dfam 4.0 family bears no similarity to any family in the column’s library. The similarity metric reflects both sequence divergence and query coverage in a single value: a low value (orange or yellow) usually indicates that the Dfam 4.0 family has no orthologous match in the column’s library, but that it has some low-scoring partial match to a non-orthologous family (for example, because both families share an ORF of common ancestry). A single entry in a contributing library can therefore account for several cells in its column: one for its true counterpart in Dfam 4.0, and others for related Dfam 4.0 families that the library does not represent. We found that the old Repbase models grouped under “Unknown” (asterisk) did not represent TEs; they were accidentally left in Dfam 4.0 but will be removed from Dfam 4.1. Excluded from this figure are: ARTEFACTS, rRNA models, and the SINE element LmeSINE1c found in the coelachanth but erroneously assigned to the Metazoa taxonomic level in Dfam.

**Figure 2.**
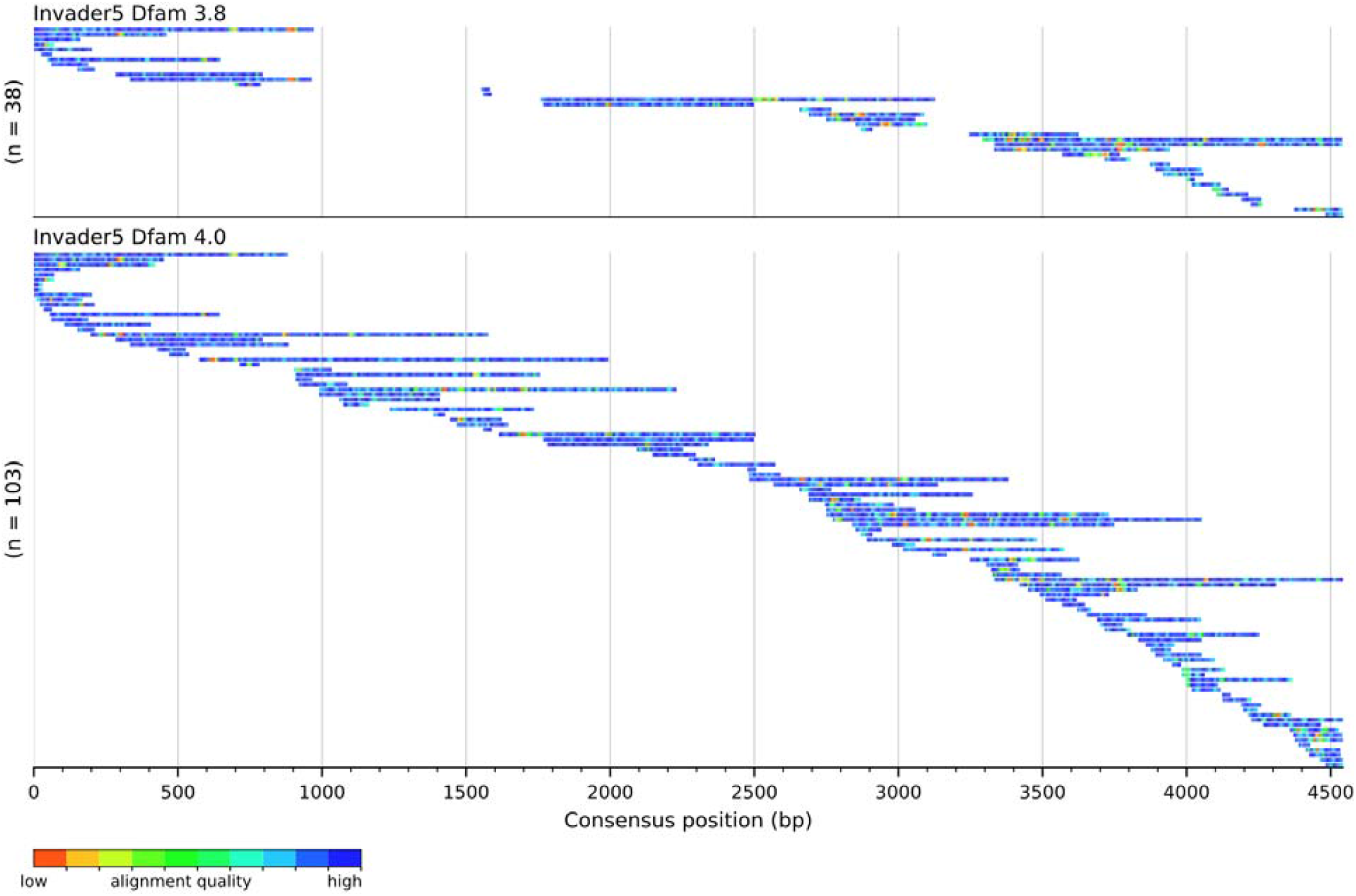
Seed alignments for the internal sequence of the Gypsy element Invader5 in Dfam 3.8 (top) and Dfam 4.0 (bottom). Each row represents an alignment, where the color range indicates divergence (orange to blue: low to high similarity). This element was active before the speciation of *D. melanogaster* from *D. simulans* and relatives. Its copies are highly fragmented. In Dfam 3.8 the alignment was biased by a single copy with large deletions in it. For the new alignment we added copies found in the genomes of *D. simulans*, *D. sechellia* and *D. yakuba*, enabling us to reconstruct a complete internal sequence with intact open reading frames.

Indeed, several families present in Repbase were absent in Dfam 3.8, due to the lack of copies in dm6 supporting the creation of a TE family model (Figure 1). A textbook example is the P element (P_Dmel); it is ubiquitous among wild *D. melanogaster* populations, but absent from dm6 and was absent from the Repbase release used for the inception of Dfam^17^, so no Dfam model existed for it until now. The new Dfam model for P_Dmel (DF003894184) is now well supported throughout the consensus by copies present amongst the 13 genomes of ^10^.

Several other specific changes in Dfam 4.0 are worth reporting. Domesticated telomeric elements, which are essential for maintaining the telomere size in *D. melanogaster*, remained challenging to curate: TART_B1 for instance, had a very poor seed alignment that could not be fixed with our augmented assembly. Instead, the updated model uses the single sequence DMU14101 (Genbank U14101.1) in order to maintain the sequence of the functional TE in the database. In addition, the subtelomeric satellite HETRP_DM (formerly DF000001643), which previously contained unspecified nucleotides (N) across 25% of its length, was removed from the database, along with TLD2 (formerly DF000001717) which is in fact a sub-fragment of Gypsy-8_DMel-I (DF003894185). Of note, the main family of rolling-circle (RC) elements (Class II), DNAREP1_DM (DF000001586) was reclassified from Helitron to Helentron after manual curation.

### Impact on overall annotation

Reannotation of the reference dm6 genome with the updated library yields an additional 2.1 Mb of repeats, or 1.55% of the genome (only considering the non-N bases on the main chromosomes, Table 1). The modest gain is due to the fact that most new or improved models describe low copy number TEs. In this annotation, the dm6 genome includes 16.47% TEs and 3.46% other repeats (satellites, simple_repeats, low_complexity, ARTEFACT and rRNA; Figure 3A,B).

**Figure 3.**
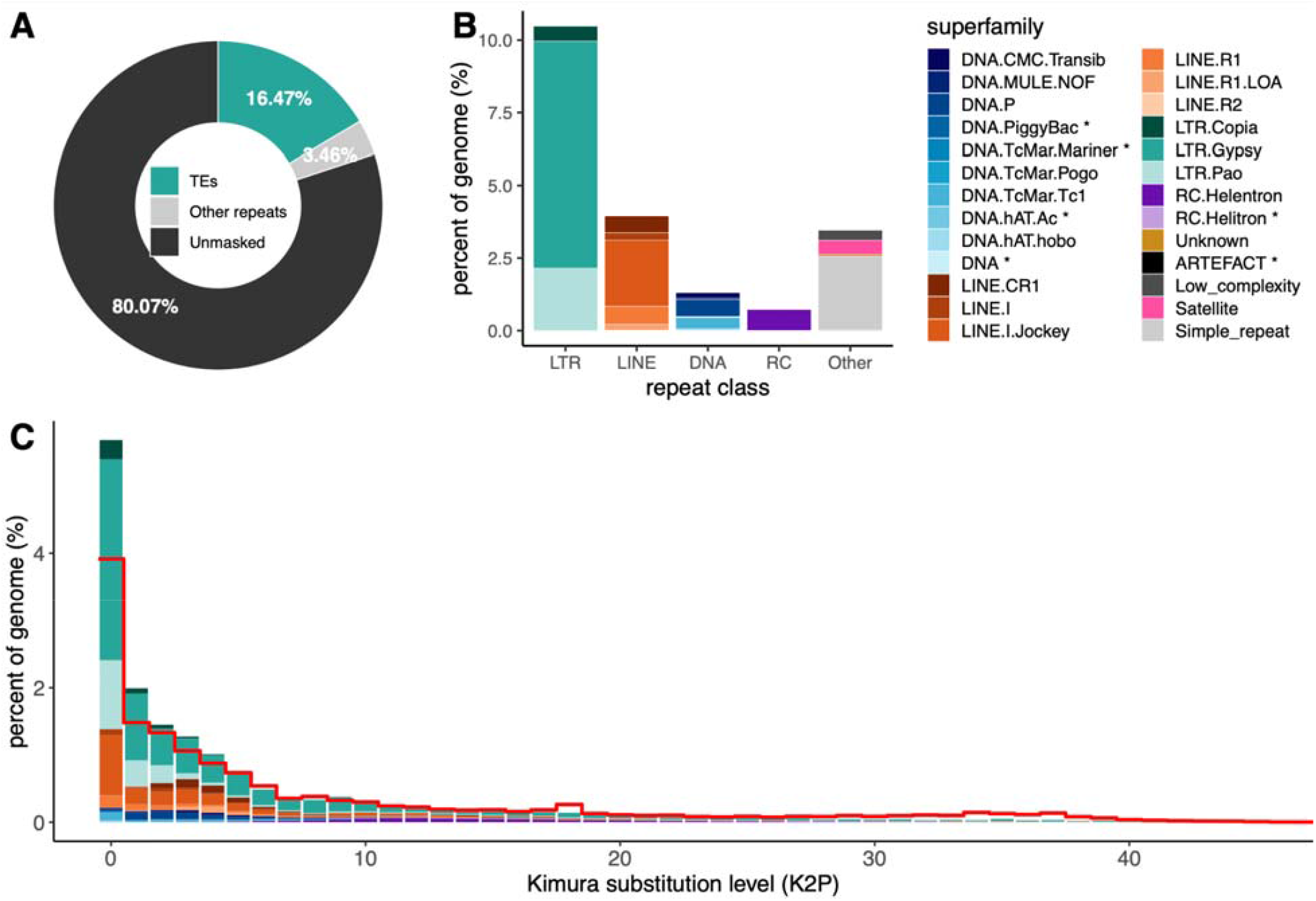
Transposable Elements annotation of the *D. melanogaster* reference genome dm6 by RepeatMasker and Dfam4.0. Chromosomes 2L, 2R, 3L, 3R, 4, X, Y and M only. Percentages exclude “N” nucleotides. **A.** Estimation of the total TE and other repeat content by RepeatMasker. **B.** Breakdown per TE subfamilies and other repeat classes. Categories with an asterisk (*) appended represent less than 0.01% of the genome. **C.** Transposable Element landscape representing binned Kimura substitution between TE copies and their consensus sequence in the library as a proxy of their evolutionary age. Filled colored bars represent annotations from Dfam 4.0; the red line marks the landscape footprint (per-bin maximum) according to Dfam3.8. Repeats in the “Other” category are excluded from the landscape.

**Table 1.** Coverage gain between Dfam3.8 and Dfam4.0 for *D. melanogaster* libraries. Reference genome dm6, chromosomes 2L, 2R, 3L, 3R, 4, X, Y and M only. Counts and percentages exclude “N” nucleotides.

| Genome size<br>(bp, no Ns) | Dfam<br>version | Interspersed repeats<br>(bp) | All repeats<br>(bp) | Increase<br>(bp) | Increase<br>(% of genome) |
| --- | --- | --- | --- | --- | --- |
| 137,077,112 | 3.8 | 20,441,791 | 25,196,857 |  |  |
|  | 4.0 | 22,673,436 | 27,321,178 | 2,124,321 | 1.55% |

A TE landscape represents the distribution of the divergence between annotated copies and their consensus, and approximates the relative age of TE families within the genome assembly. An updated TE landscape shows a sharp skew to the left (towards newer TE families) in comparison to Dfam 3.8 (Figure 3C). This was expected after the refinement of the models, because the new analysis included representative seeds across multiple genomes, tightening the apparent divergence spread within a family, despite the addition of a number of older elements predating the speciation of the *D. melanogaster* species subgroup. This updated TE landscape reinforces the established paradigm in *D. melanogaster* of a highly dynamic repeatome, in which the activity of numerous TE families is counterbalanced by a high deletion rate and strong purifying selection.

### Feedback and future revisions

Dfam is a collaborative platform. Error correction, feedback, new entries are welcome. Questions and remarks can be addressed at.

## Ethics approval and consent to participate

Not applicable

## Consent for publication

Not applicable

## Availability of data and materials

The Dfam 4.0 libraries are available online at www.dfam.org; documentation to extract Dfam libraries are available at https://dfam.org/releases/current/families/README.txt and requests for help can be sent at. Scripts and wrappers used for this work are available online at https://github.com/Dfam-consortium/dfam40-dmel-paper.

## Author’s contribution

RH, AS, and CG conceptualized the project. RH, AS and CG performed the experiments and conducted bioinformatic analyses. AS created the TE models. RH populated and deployed the database. TJW and AS supervised the project. All authors contributed to the writing of the manuscript.

## Acknowledgements

The authors would like to thank Gabriel Rech, Josefa Gonzalez, Riccardo Pianezza and Sarah Signor for their contribution to the Dfam database.

## Footnotes

* We note that a subsequent version 10.1, last updated Aug. 27 2021 by the Bergman lab is available^36^

## References

1. Jennings, B. H. Drosophila – a versatile model in biology & medicine. Mater. Today (Kidlington) 14, 190–195 (2011).

2. Morgan, T. H. Sex limited inheritance in Drosophila. Science 32, 120–122 (1910).

3. Green, M. M. Transposable elements in Drosophila and other Diptera. Annu. Rev. Genet. 14, 109– 120 (1980).

4. Petrov, D. A., Lozovskaya, E. R. & Hartl, D. L. High intrinsic rate of DNA loss in Drosophila. Nature 384, 346–349 (1996).

5. Petrov, D. A., Fiston-Lavier, A.-S., Lipatov, M., Lenkov, K. & González, J. Population genomics of transposable elements in Drosophila melanogaster. Mol. Biol. Evol. 28, 1633–1644 (2011).

6. Charlesworth, B. & Langley, C. H. The population genetics of Drosophila transposable elements. Annu. Rev. Genet. 23, 251–287 (1989).

7. Mérel, V., Boulesteix, M., Fablet, M. & Vieira, C. Transposable elements in Drosophila. Mob. DNA 11, 23 (2020).

8. Kapun, M. et al. Genomic analysis of European Drosophila melanogaster populations reveals longitudinal structure, continent-wide selection, and previously unknown DNA viruses. Mol. Biol. Evol. 37, 2661–2678 (2020).

9. Lerat, E. et al. Population-specific dynamics and selection patterns of transposable element insertions in European natural populations. Mol. Ecol. 28, 1506–1522 (2019).

10. Rech, G. E. et al. Population-scale long-read sequencing uncovers transposable elements associated with gene expression variation and adaptive signatures in Drosophila. Nat. Commun. 13, 1948 (2022).

11. Rubin, G. M., Kidwell, M. G. & Bingham, P. M. The molecular basis of P-M hybrid dysgenesis: the nature of induced mutations. Cell 29, 987–994 (1982).

12. Pianezza, R., Scarpa, A., Haider, A., Signor, S. & Kofler, R. Spatiotemporal tracking of three novel transposable element invasions in Drosophila melanogaster over the last 30 years. Mol. Biol. Evol. 42, msaf143 (2025).

13. Scarpa, A. et al. Double trouble: two retrotransposons triggered a cascade of invasions in Drosophila species within the last 50 years. Nat. Commun. 16, 516 (2025).

14. Bao, W., Kojima, K. K. & Kohany, O. Repbase Update, a database of repetitive elements in eukaryotic genomes. Mob. DNA 6, 11 (2015).

15. The Berkeley Drosophila Genome Project. fruitfly.org https://www.fruitfly.org/.

16. Öztürk-Çolak, A. et al. FlyBase: updates to the Drosophila genes and genomes database. Genetics 227, iyad211 (2024).

17. Hubley, R. et al. The Dfam database of repetitive DNA families. Nucleic Acids Res. 44, D81–9 (2016).

18. Bargues, N. & Lerat, E. Evolutionary history of LTR-retrotransposons among 20 Drosophila species. Mob. DNA 8, 7 (2017).

19. Smit, A. F., Tóth, G., Riggs, A. D. & Jurka, J. Ancestral, mammalian-wide subfamilies of LINE-1 repetitive sequences. J. Mol. Biol. 246, 401–417 (1995).

20. Smit, A. F. Identification of a new, abundant superfamily of mammalian LTR-transposons. Nucleic Acids Res. 21, 1863–1872 (1993).

21. Hubley, R., & Smit, A. F. A. COSEG: A Program to Identify Repeat Subfamilies Using Significant Co-Segregating Mutations. (2008–2026).

23. Storer, J. M., Hubley, R., Rosen, J. & Smit, A. F. A. Curation guidelines for de novo generated transposable element families. Curr. Protoc. 1, e154 (2021).

23. Kohany, O., Gentles, A. J., Hankus, L. & Jurka, J. Annotation, submission and screening of repetitive elements in Repbase: RepbaseSubmitter and Censor. BMC Bioinformatics 7, 474 (2006).

24. Quesneville, H. et al. Combined evidence annotation of transposable elements in genome sequences. PLoS Comput. Biol. 1, 166–175 (2005).

25. Carey, K. M., Patterson, G. & Wheeler, T. J. Transposable element subfamily annotation has a reproducibility problem. Mob. DNA 12, 4 (2021).

27. Berkeley Drosophila Genome Project, Berkeley (CA). Natural transposable element dataset, version 9.4.1. (2008).

27. Evgen’ev, M. B. et al. Penelope, a new family of transposable elements and its possible role in hybrid dysgenesis in Drosophila virilis. Proc. Natl. Acad. Sci. U. S. A. 94, 196–201 (1997).

28. Pyatkov, K. I. et al. Penelope retroelements from Drosophila virilis are active after transformation of Drosophila melanogaster. Proc. Natl. Acad. Sci. U. S. A. 99, 16150–16155 (2002).

29. Suvorov, A. et al. Widespread introgression across a phylogeny of 155 Drosophila genomes. Curr. Biol. 32, 111–123.e5 (2022).

30. Chan, P. P. & Lowe, T. M. GtRNAdb 2.0: an expanded database of transfer RNA genes identified in complete and draft genomes. Nucleic Acids Res. 44, D184–9 (2016).

31. Groza, C., Chen, X., Wheeler, T., Bourque, G. & Goubert, C. A unified framework to analyze transposable element insertion polymorphisms using graph genomes. Nature Communications 15, 8915 (2024).

32. Heller, D. & Vingron, M. SVIM-asm: structural variant detection from haploid and diploid genome assemblies. Bioinformatics 36, 5519–5521 (2021).

33. Li, H. Minimap2: pairwise alignment for nucleotide sequences. Bioinformatics 34, 3094–3100 (2018).

34. Flynn, J. M. et al. RepeatModeler2 for automated genomic discovery of transposable element families. Proc. Natl. Acad. Sci. U. S. A. 117, 9451–9457 (2020).

35. Perez, G. et al. The UCSC Genome Browser database: 2025 update. Nucleic Acids Res. 53, D1243– D1249 (2025).

36. Bergman, C. Drosophila transposable element canonical sequences. https://github.com/bergmanlab/drosophila-transposons (2021).

